# CryoEM insights into RNA primer synthesis by the human primosome

**DOI:** 10.1101/2023.07.20.549859

**Authors:** Zhan Yin, Mairi L. Kilkenny, De-Sheng Ker, Luca Pellegrini

## Abstract

Eukaryotic DNA replication depends on the primosome – a complex of DNA polymerase alpha (Pol α) and primase – to initiate DNA synthesis by polymerisation of an RNA - DNA primer. Primer synthesis requires the tight coordination of primase and polymerase activities. Recent cryo-electron microscopy (cryoEM) analyses have elucidated the extensive conformational transitions required for RNA primer handover between primase and Pol α and primer elongation by Pol α. Because of the intrinsic flexibility of the primosome however, structural information about the initiation of RNA primer synthesis is still lacking. Here, we capture cryoEM snapshots of the priming reaction to reveal the conformational trajectory of the human primosome that brings the PRIM1 and PRIM2 subunits of primase together, poised for RNA synthesis. Furthermore, we provide experimental evidence for the continuous association of primase subunit PRIM2 with the RNA primer during primer synthesis and for how both initiation and termination of RNA primer polymerisation are licensed by specific rearrangements of POLA1, the polymerase subunit of Pol α. Our findings fill a critical gap in our understanding of the conformational changes that underpin the synthesis of the RNA primer by the primosome. Together with existing evidence, they provide a complete description of the structural dynamics of the human primosome during DNA replication initiation.

## INTRODUCTION

Duplication of the genome before cell division is performed by the replisome, a large multi-protein assembly equipped with the necessary enzymatic activities for DNA replication^1^. In all kingdoms of life, replisomes solve the problem of initiating chromosomal DNA synthesis by the intervention of a specialised DNA-dependent RNA polymerase known as primase^2^. In the eukaryotic replisome, a heterotetrameric complex of DNA polymerase alpha (Pol α) and primase known as primosome is responsible for priming DNA synthesis^3^. The primosome initiates nucleotide polymerisation by assembling an RNA - DNA primer that is extended by DNA polymerases ε and δ on the leading and lagging strands, respectively^4^. Due to the semi-discontinuous nature of DNA replication, primosome activity is constantly required for Okazaki fragment production on the lagging-strand template^5^. In addition to its critical replicative role, the human primosome participates in the processes of fill-in synthesis during telomere maintenance^6^ and repair of DNA double-strand breaks^7^. Partial deficiency in POLA1, the catalytic subunit of Pol α, is the causative agent of genetic conditions such as X-linked reticulate pigmentary disorder and van Esch – O’ Driscoll syndrome, characterised by immunological and developmental defects^8^.

The archaeo-eukaryotic primase is a heterodimer of catalytic (PRIM1) and regulatory (PRIM2) subunits^9–11^. PRIM1’s active site has a simple architecture made of an open groove formed by two adjoining β-sheets and lined by aspartate residues that perform metal-ion dependent catalysis^10, 12^. PRIM2 consists of an N-terminal domain (PRIM2_NTD_) that docks onto the side of PRIM1 and is flexibly linked to the C-terminal domain (PRIM2_CTD_) that can reach over PRIM1’s active site. The PRIM2_CTD_ harbours residues that are important for catalysis and facilitates initiation and elongation of the RNA primer^13, 14^, while the span of its flexible linker help define the size of the RNA primer^15^. Pol α shares an evolutionarily conserved subunit architecture with Pol ε and δ, consisting of a catalytic subunit POLA1 that is constitutively bound via a large C-terminal domain (POLA1_CTD_) to its accessory B subunit (POLA2)^16, 17^. Both POLA1 and POLA2 contain largely unstructured N-terminal extensions that mediate protein interactions and are not involved in primer synthesis^18–20^. A short motif at the POLA1 C-end tethers Pol α to primase and is essential for holding them together in the primosome^21^.

Primer synthesis by the primosome requires the coordination of its primase and polymerase activities. The PRIM1 and PRIM2 subunits of primase cooperate to initiate and extend an RNA primer to about 7 – 12 nucleotides; the primer is then transferred intramolecularly to the active site of POLA1 for deoxynucleotide extension, yielding an RNA-DNA primer of 20-30 nucleotides^22–24^. Execution of this series of steps by the primosome requires a complicated series of conformational transitions. Such complexity has made the structural analysis of primer synthesis a highly challenging task. Early low-resolution EM studies showed that the yeast primosome is organised as two loosely connected primase and polymerase lobes^14^. Surprisingly, a crystal structure of the human primosome revealed that, in the absence of template DNA and nucleotides, the structure adopts a specific conformation incompatible with primer synthesis^25^. This self-inhibited conformation was later confirmed by cryoEM^26^. In this apo state, PRIM1 and PRIM2_CTD_ are held widely apart by multiple interactions with the polymerase and carboxy-terminal domains of POLA1, while POLA2 blocks access to the primer-template binding site of the POLA1 polymerase domain^25^.

A recently reported cryoEM of the human primosome in the presence of an RNA primer and DNA template captured the enzyme in a *bona fide* elongation state, showing how the 3’-end extension of the primer with deoxynucleotides by POLA1 can take place concurrently with continuous binding of the PRIM2_CTD_ to the nucleotide triphosphate at the 5’-end of the primer^27^. Structures of Xenopus and yeast primosomes provided with primed templates have confirmed the elongation conformation of the enzyme and proposed mechanistic models for the transfer of the mature RNA primer to POLA1 for deoxynucleotide extension^28, 29^.

Structural work also has explored the interaction of the telomeric CST (CTC1-STN1-TEN1) complex with Pol α - primase. CryoEM structures of CST bound to the apo state of the human and Tetrahymena primosomes have shown how the CST can recruit the primosome via interactions of CST’s large CTC1 subunit with PRIM2_NTD_ and POLA1_CTD_^30, 31^. Further cryoEM structures in the presence of telomeric DNA have revealed the extensive primosome remodelling induced by the CST, which binds the telomeric repeat template and feeds it to the primase subunits, held in the appropriate relative position for initiation^32^.

A common theme highlighted by these primosome structures is the extreme flexibility of the enzyme and the large rearrangements that it must undergo during primer synthesis. Interestingly, the apo state of the yeast primosome has revealed a more loosely connected PRIM1 subunit than observed in the apo human primosome^28^; furthermore, the purified primosome from *Xenopus laevis* is present as a mixture of apo state and a more open form where the POLA1 polymerase domain and PRIM2_CTD_ appear largely disordered^29^. In contrast, the apo state of the human primosome consists exclusively of its self-inhibited conformation, which even CST binding doesn’t substantially alter^30^, suggesting a different degree of primosome regulation among species.

The important mechanistic insights delivered by the available primosome structures were obtained by providing the enzyme with pre-formed primer: template substrates ^24–26^. These structures therefore do not inform us about the critical first step of primer synthesis, when the primosome transitions from the apo state to one that is competent for initiation. Here we used cryoEM of priming reactions snap-frozen at different times and conditions to obtain structural intermediates of the human primosome, that let us describe its trajectory from the apo state to the initiation stage. We further provide evidence that supports the continuous association of the PRIM2_CTD_ with the 5’-end of the RNA primer postulated by current models of priming and show that RNA primer synthesis is tightly coupled to rearrangements of the POLA1 polymerase domain.

## MATERIALS AND METHODS

### Primosome cloning and expression

The human primosome (POLA1: 334-1462; POLA2:149- 598; PRIM1: 1-420; PRIM2: 1-509) was expressed in Sf9 insect cells and purified as described previously^19^. In brief, the pFBDM vector was employed for cloning the double StrepII-tagged human POLA1 and POLA2 subunits, while a separate pFBDM vector was used for cloning the His_10_-tagged human PRIM1 and PRIM2. Both vectors were then utilized to create a recombinant baculovirus, which was co-infected into Sf9 insect cells for expression. Following a 72-hour incubation at 27°C, the cells were harvested.

### Protein purification

Sf9 cells were lysed by sonication with 100 mL lysate buffer containing 20 mM Tris-HCl pH 7.4, 300 mM KCl, 1 mM TCEP, 10 μL of 25 U/μL Benzonase, 1 mg bovine pancreas RNaseA and EDTA-free Protease Inhibitor Cocktail (Roche). Then the cell suspension was centrifuged at 45,000 g for 1.5 hours. The supernatant was filtered through a 5.0 µm syringe filter followed by 0.45 µm syringe filters and loaded on to 3 mL gravity flow column packed with StrepTactin Sepharose resin (IBA Lifesciences). The column was washed with wash buffer (20 mM Tris-HCl pH 7.4, 300 mM KCl, 1mM TCEP) and elute with elution buffer (20 mM Tris-HCl pH 7.4, 300 mM KCl, 1 mM TCEP, 7.5 mM d-Desthiobiotin). The elution fractions were then loaded onto ÄKTA purifier system (GE Healthcare) connected with 5mL HisTrap HP column (Cytiva). The sample was washed with HisTrap wash buffer (20 mM Tris-HCl pH 7.4, 300 mM KCl, 40 mM imidazole) and eluted in the elution buffer (20 mM Tris-HCl pH 7.4, 300 mM KCl, 250 mM imidazole). The elution fractions were collected, concentrated and buffer exchanged in 20 mM Tris-HCl pH 7.4, 300 mM KCl by using Vivaspin 500 MWCO 100 000 Centrifugal Concentrator (Sartorius). The human primosome was snap-frozen in liquid nitrogen and stored at –80 °C.

### Primosome activity assays

The gel-based primosome activity assays, which had been previously described^33^, underwent modifications and adjustments for the purpose of this study. Specifically, the reaction buffer contains 25 mM HEPES pH 7.4, 120 mM NaCl, 1 mM DTT, 10 mM Mg(OAc)_2_ and 2 mM Mn(OAc)_2_. For primosome initiation assays, the reaction buffer was combined with 5 µM PolydT-70mer ssDNA template (IDT), 0.5 mM ATP, and 0.1 µM human primosome. The resulting mixture was incubated at 20 °C for a duration, depending on the specific experimental conditions. For RNA primer handover assay, dATP and Cy5-dATP (Stratech) was added to the reaction when indicated. The reaction was terminated by adding loading buffer (95% formamide, 25 mM EDTA and 0.01% xylene cyanol and bromophenol blue) and heated to 95°C for 5 mins. Samples were loaded onto 19% urea-polyacrylamide gel and run in 1x TBE buffer at 500V for 2.5 hours. Then the gel was stained with Sybr Gold Stain (Thermo Fisher Scientific) and visualised by scanning with Typhoon FLA 9000 (GE Healthcare).

### Cryo-EM sample preparation and grid freezing

CryoEM samples were optimised to match the primosome reaction conditions. Purified human primosome was incubated with 5 µM PolydT-70mer ssDNA template, 0.5 mM ATP in reaction buffer (25 mM HEPES pH 7.4, 120 mM NaCl, 1 mM TECP, 10 mM Mg(OAc)_2_ and 2 mM Mn(OAc)_2_) for 5 mins at 20 °C. The samples for elongation and handover were made in the same reaction condition with additional of 0.5 mM dATP. A mild crosslinking of 1 mM BS^3^ was applied to the samples after stopping the reaction on ice for 30 mins. UltrAuFoil R1.2/1.3 or R0.6/1 300 mesh gold grids (Quantifoil Micro Tools GmbH) were glow discharged on both sides for 1.5 mins on a PELCO easiGlow glow discharge unit (20 mA, 0.4 mbar). Human primosome under reaction conditions was applied on grids using an FEI Vitrobot Mark IV (FEI), set to 4 °C in 100% humidity, 10 s blotting time, –5 blotting force.

### Cryo-EM data collection, image processing and model building

The highest quality cryo-frozen grids were identified using a 200 keV Talos Arctica transmission electron microscope (Thermo Fisher Scientific), then high-resolution data acquisition was performed on a 300 keV Titan Krios microscope (Thermo Fisher Scientific) fitted with a Gatan K3 detector with AFIS mode at the Cambridge University cryo-EM facility in the Department of Biochemistry. Four data sets were collected with different reaction conditions. All the micrographs are collected at a defocus range from −1 to −2.6 μm; and the calibrated pixel size was 1.066 Å.

A total of 15,430 micrographs were collected for primosome initiation with ssDNA, ATP and dATP. Each micrograph was collected with a total dose of 47.78 e^−^/Å^2^ in 45 frames. The micrographs were motion-corrected and CTF fitted in Warp and the particles are initially picked with pre-trained BoxNet model in Warp with a box size of 380 pixels^34^. Then the picked 1,542,848 particles were imported in CryoSPARC v4.0.0^35^ and first three rounds of 2D classification were to remove non-protein junk. The resulting 714,412 particles went through ab-initio in 5 classes. Particles from two intact ab-initio models were selected and the non-uniform refinements were performed, resulting in an initiation map showing the platform and a pre-initiation stage 1 map with a global resolution of 3.07 Å based on the 0.143 FSC criterion.

A total of 7,014 micrographs were collected for primosome with ssDNA, ATP, dATP and BS^3^ crosslinking. Each micrograph was collected with a total dose of 50.50 e^−^/Å^2^ in 45 frames, using the FEI EPU software. The micrographs were motion-corrected and CTF fitted in Relion 4.0^36^. Then the motion-corrected images were imported into CryoSPARC v4.2.1 and two rounds of 2D classification were preformed to remove bad particles, resulting in 1,122,827 good particles. Particles were then separated into subsets for another round of 2D classification. 2D classes showing a clear apo-like feature were selected and went through ab-initio in 5 classes. Two groups of particles showing intact ab-initio models were selected and further subclassified through ab-initio and non-uniform refinements. The map refined from 132,036 particles went through 3D variability analysis, and particles are further clustered into 3 subclasses, and the movement of flexible subunits was recorded in a movie. The 3 subclasses were then further non-uniform refined and resulted in 3 pre-initiation maps (pre-initiation stage 2,3 and 4) with a global resolution of 4.11 Å, 3.97 Å, 4.21 Å respectively based on the 0143 FSC criterion. The particles from 2D classes without apo-like feature went through two runs of ab-initio to removed junk particles. The selected 53,489 particles were further non-uniform refined, resulting in an initiation stage map (initiation stage I) with PRIM2_CTD_ bound with a global resolution of 6 Å based on the 0.143 FSC criterion.

A total of 10,677 micrographs were collected for primosome with ssDNA and ATP. Each micrograph was collected with a total dose of 46.05 e^−^/Å^2^ in 40 frames. The micrographs were motion-corrected and CTF fitted in Warp and the particles are initially picked with pre-trained BoxNet model in Warp with a box size of 380 pixels^34^. Four rounds of 2D classification were performed to remove the non-protein junk. 510,178 particles went through three rounds of abinitio reconstructions due to high heterogeneity. And a class of 56,798 particles were selected for non-uniform refinement, resulting in an initiation stage map (initiation II) with a global resolution of 4.03 Å based on the 0.143 FSC criterion. A further 3D viability analysis was performed and the movement of PRIM2_CTD_ was recorded in a movie.

A total of 14,621 micrographs were collected for primosome with ssDNA ATP, dATP and BS^3^ crosslinking with a longer reaction time. Each micrograph was collected with a total dose of 53.67 e^−^/Å^2^ in 45 frames, using the FEI EPU software. The micrographs were motion-corrected and CTF fitted in Relion 4.0^36^. Then the motion-corrected images were imported into CryoSPARC v4.2.1 and particles were selected by topaz trained with the manually selected reference particles from a subset of 500 micrographs^37^. Four rounds of 2D classification were preformed to remove bad particles, resulting in 374,350 good particles. Then the good particles went through ab-initio in 2 classes. One of the ab-initio models with 196,320 particles went through non-uniform refinement showing a map at initiation stage with a global resolution of 3.73 Å; Another set of 178,030 particles went through non-uniform refinement showing a map (handover state) with a global resolution of 6.15 Å based on the 0.143 FSC criterion.

Three models from PDB, 7OPL, 5EXR and 6R4S, were used as initial models, and all the maps were rigid body-fitted in UCSF Chimera X v1.6. In pre-initiation stage 2 to 4, PRIM2_CTD_ was fitted by fit-in-map function of ChimeraX, and a local likelihood-based docking was performed by using localEM fitting in ChimeraX^38^.

## RESULTS

In this study, we sought to exploit cryoEM’s power to identify distinct structural intermediates in a sample to capture the primosome during the critical steps of RNA primer initiation and elongation. To this aim, we performed priming reactions containing purified primosome, poly-dT (70 nts) as template DNA and either ATP alone or ATP and dATP, which were directly frozen on grids at appropriate reaction times for subsequent cryoEM analysis (Figure 1a, 1b). The reaction conditions were optimised by urea-PAGE analysis of reaction products (Figure S1). Using this approach, we succeeded in capturing several structural intermediates of the primosome that shed light on its conformational rearrangements as it transits from the initial inactive state to initiation of RNA primer synthesis and primer handover to POLA1.

**Figure 1.**
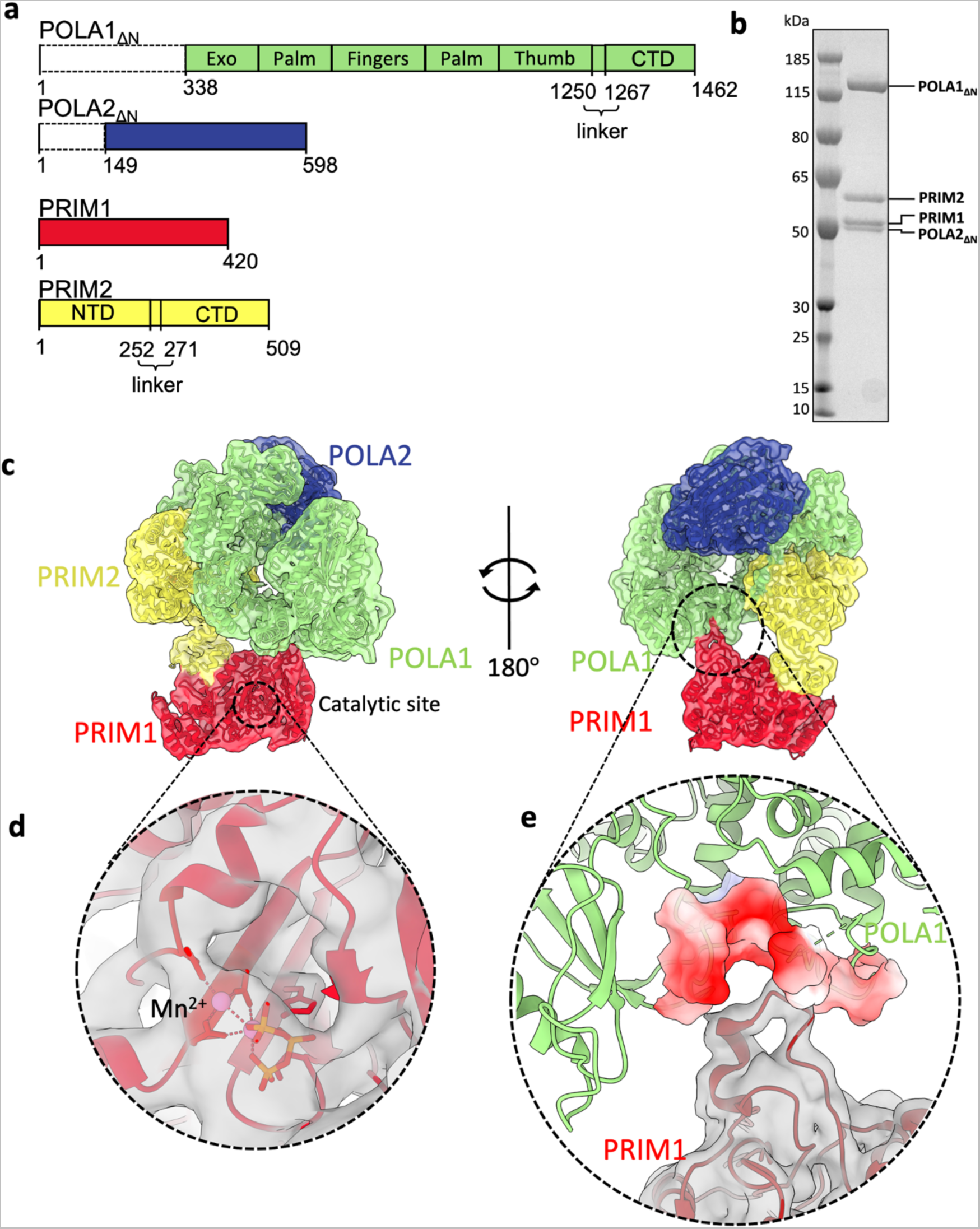
Structure of the human primosome in the pre-initiation stage 1. (**a**) Subunit composition and domain structure of the human primosome. The extended, largely disordered N-terminal sequences (dashed boxes) have been omitted in the recombinant POLA1 and POLA2 proteins used in this study. (**b**) SDS-PAGE of purified human primosome. (**c**) Structures for primosome (PDB: 7OPL) and PRIM1 (PDB: 6R4S) were rigid-body fitted into the cryoEM map (EMD-17795) of pre-initiation stage 1. (**d**) Density for a nucleotide was identified at the catalytic site of PRIM1. (**e**) PRIM1 loop N92 - Q100 is docked in a negatively charged groove between the finger and palm of POLA1.

### Unlocking the auto-inhibitory state of the human primosome

Our protocol of sample preparation purposedly omitted any device to enrich for a specific primosome intermediate (Table S1, Figure S2). Consequently, identification of structural intermediates in our reaction samples required extensive 2D- and 3D-classification. The earliest primosome intermediate (pre-initiation stage 1) that we could identify in a reaction with both ATP and dATP corresponded to the previously described ‘apo’ conformation, with additional nucleotide density at the catalytic site of PRIM1 (Figure 1c, 1d, Figure S3). In this self-inhibited apo state, the human primosome adopts a compact configuration where PRIM1 and PRIM2_CTD_ are held wide apart by extensive interactions with the polymerase and carboxy-terminal domains of POLA1, while POLA2 sits on top of POLA1 polymerase domain, blocking access of the primer/template duplex to the active site. In this conformation, PRIM1 is sequestered away from the PRIM2_CTD_ by an interaction of its ‘anchor’ loop formed by residues N92 to Q100 with POLA1. The anchor loop protrudes from the perimeter of PRIM1’s primase fold to dock in a negatively-charged groove at the junction of finger and palm in POLA1 polymerase domain (Figure 1e).

Our pre-initiation stage 1 structure shows that nucleotide binding by PRIM1 does not induce large-scale conformational changes in the human primosome, indicating that its presence is not sufficient to unlock the auto-inhibitory state. To determine whether template DNA can induce the required changes in primosome structure, we incubated the primosome with ssDNA without nucleotides for cryoEM analysis. Extensive 2D classification of these primosome particles revealed only the presence of the apo form (Figure S4a), indicating that – as in the case of the nucleotide – binding of the ssDNA template alone is not able to trigger conformational changes in the human primosome. Accordingly, activity studies showed that pre-incubation with either nucleotide or ssDNA does not accelerate the reaction (Figure S4b), which agrees with our structural observations.

### The primosome trajectory from self-inhibited state to initiation state

To capture the transition from apo to initiation state, we repeated the cryoEM analysis of the initiation reaction with samples that had been subjected to mild crosslinking and were able to detect additional conformational states of the primosome that had not been initially apparent (Table S1, Figure S5). We could identify three further pre-initiation states that we refer to as stages 2 to 4. In these conformations, the anchor loop of PRIM1 has been released from its dock on POLA1, allowing PRIM1 to move up along the thumb of POLA1 in a rigid body motion bringing it closer to the PRIM2_CTD_ (Figure 2a, Movie S1). PRIM1’s rigid-body shift from stage 1 to stage 2 was the largest at over 20 Å, followed by a further shift of under 10 Å in the same direction (stage 3). PRIM2_CTD_ moved in concert with PRIM1, sliding down POLA1’s thumb domain and getting progressively closer to PRIM1 in stages 2 and 3 (Figure 2b).

**Figure 2.**
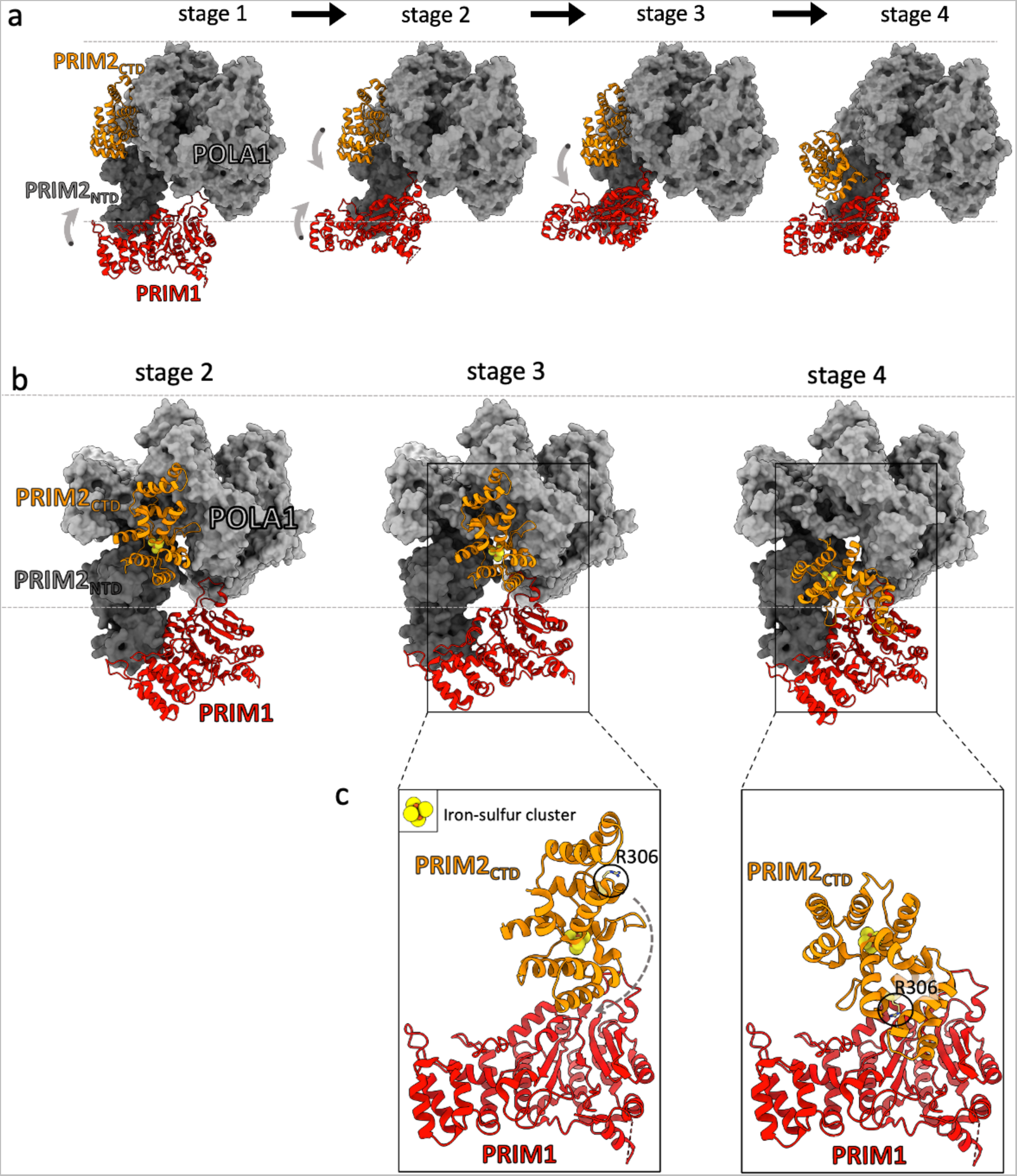
The concerted movements of primase subunit PRIM1 and PRIM2 towards each other in the pre-initiation states. (**a**) Primosome structures at the four pre-initiation stages. PRIM1 and PRIM2_CTD_ are shown in red and orange ribbons, respectively, the rest of the primosome is shown as molecular surface in grey. (**b**) Side-view of PRIM2_CTD_ movement from stage 2 to 4. (**c**) Between stages 3 and 4, PRIM2_CTD_ undergoes a large rotation to adopt the correct position for initiation of primer synthesis. The iron-sulphur cluster and initiation residue R306 in PRIM2_CTD_ are shown.

The last stage in the transition from the self-inhibited conformation to the initiation state showed an additional large drop in the position of the PRIM2_CTD_, which brought it into contact with PRIM1 (stage 4). No further conformational stages were identified, and we therefore assume that – by stage 4 – PRIM1 and PRIM2_CTD_ are in a *bona fide* initiation conformation that is licensed for primer synthesis. Although the resolution of our stage 4 map for the PRIM2_CTD_ is insufficient to demonstrate this conclusively, the PRIM2_CTD_ must undergo a rotation of about 180° prior to docking onto PRIM1’s active site (Figure 2c); such large rearrangement is required to bring the PRIM2_CTD_ residues that participate in RNA primer synthesis, such as R306, in the correct orientation for catalysis.

### The initiation state

At pre-initiation stage 4, the polymerase domain of POLA1 is clearly visible in the map, occupying the same position as in the initial, inactive conformation of the human primosome (Figure 2a). In this conformation, POLA1’s palm domain abuts PRIM1 and PRIM2_CTD_, precluding access to the primase active site in a way that is likely to be inhibitory of primer synthesis. Further cryoEM data collection and analysis of the primosome under similar reaction conditions revealed a stage 4-like structure, with PRIM1 and PRIM2_CTD_ in the same reciprocal arrangement compatible with initiation but with missing density for the POLA1 polymerase domain (Figure 3a, Figure S5, Table S1). We suggest that this conformation represents the true initiation conformation of the primosome, where POLA1 polymerase domain has been released from its initial position while remaining tethered to primase and the POLA2 subunit via its C-terminal domain. The insufficient resolution of our map precluded the identification of the mechanism of POLA1 release, but it is likely to be triggered by a combination of nucleotide and template DNA binding, possibly coupled to local conformational changes in PRIM1 and PRIM2_CTD_.

**Figure 3.**
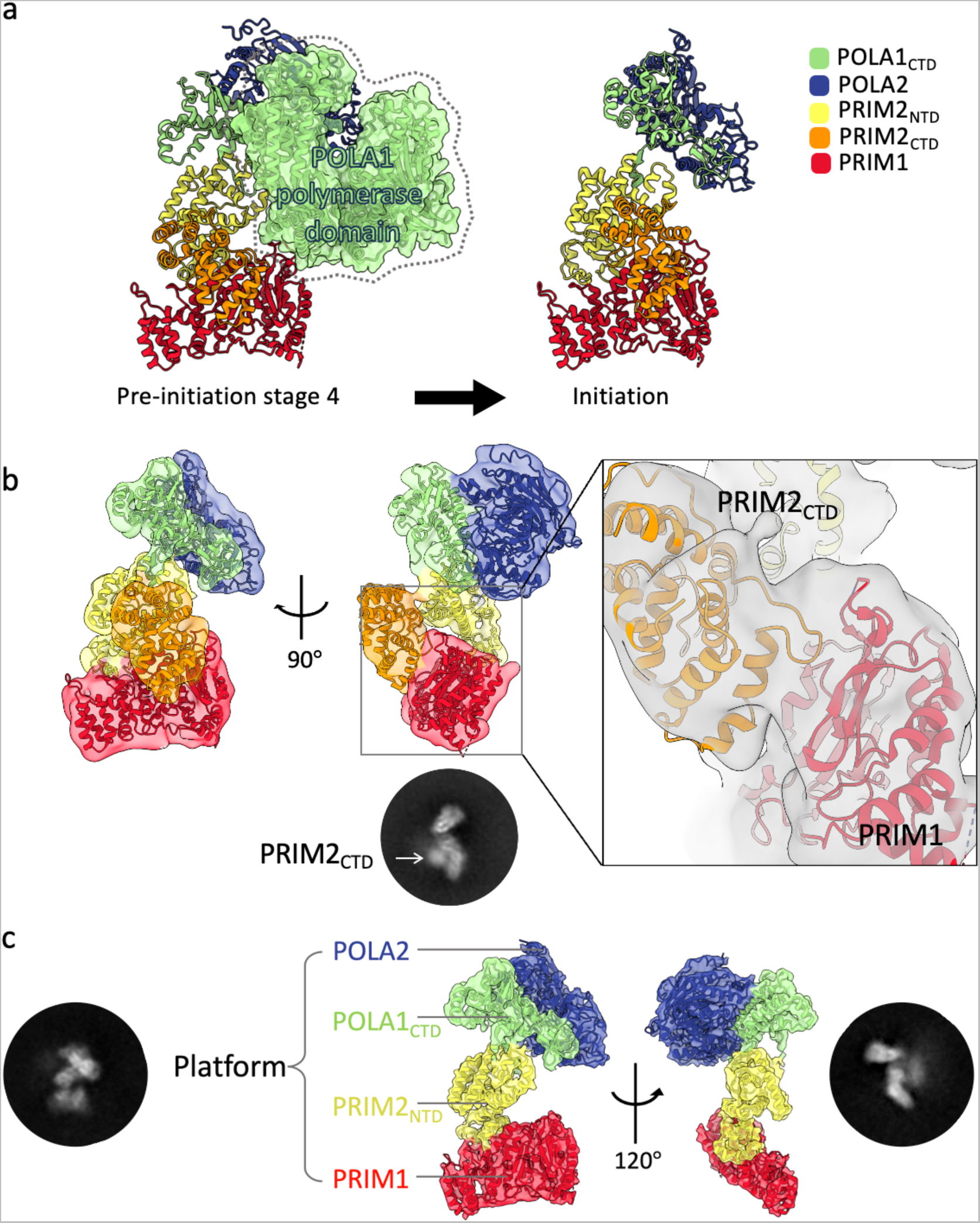
Primosome transition from pre-initiation to initiation involves release of POLA1 polymerase domain. (**a**) Primosome structures at pre-initiation stage 4 and initiation stage. The cryoEM map for POLA1 polymerase is present in pre-initiation stage 4 (shown as transparent surface) and missing at the initiation state. (**b**) CryoEM map of the initiation state (EMD-17812) with fitted primosome, showing the PRIM2_CTD_ positioned on top of PRIM1 active site, poised for RNA primer synthesis. Representative 2D class highlighting the density for PRIM2_CTD._ (**c**) CryoEM map of the primosome platform at the initiation state (EMD-17813). Primosome subunits are coloured as in (a).

In the initiation state, the release of POLA1 polymerase domain is coupled to a repositioning of POLA1_CTD_-POLA2 consisting of a rotation of about 38° coupled to a shift of 8 Å (movie S2). PRIM2_CTD_ aligns its N-terminal helix to the β-strands of PRIM1_NTD_ that harbour the active site, which becomes partially enclosed by PRIM2_CTD_ (Figure 3b). Although we cannot assign density to the DNA template, we propose that it becomes sandwiched between PRIM2_CTD_ and PRIM1, in the appropriate conformation for templating the nucleotide bound in the active site of PRIM1.

A primosome sub-assembly of PRIM1, PRIM2_NTD_, POLA1_CTD_ and POLA2 forms the ‘platform’ component of the primosome, and it remains present in the initiation state (Figure 3c). The platform has been observed in a similar conformation in the published structures of human primosome bound to RNA primer and DNA template^27^, and in the CST – primosome structure (Figure S6)^32^. Our data show that the platform persists in the different stages of RNA primer synthesis and handover, thus providing a scaffold for the conformational transitions of PRIM2_CTD_ and POLA1 polymerase domain.

### PRIM2_CTD_ tracks the growing RNA primer during synthesis

To obtain further insights into the mechanism of RNA primer synthesis, we modified our reaction conditions by using ATP only and omitting crosslinking (Table S1, Figure S7). This approach let us obtain a map of the synthesising primosome that provides insights into the conformational trajectory followed by PRIM2_CTD_ during RNA polymerisation.

Density for the POLA1 polymerase domain was not detectable in the map, confirming that it must be released from its initial position to allow for RNA primer synthesis. The PRIM2_CTD_ was also not initially visible in the map; however, inspection of the map at a lower contour threshold revealed that the PRIM2_CTD_ density had morphed from the defined volume observed above PRIM1’s active site in the initiation stage to a diffuse, elongated shape pointing away from PRIM2_CTD_’s initiation position (Figure 4a). We interpreted this feature of the map as likely to represent the movement of PRIM2_CTD_ during RNA primer synthesis. We could not isolate defined PRIM2_CTD_ intermediates in the map, indicating a continuous motion during RNA primer synthesis. Instead, we analysed the PRIM2_CTD_ density by 3D Variability Analysis in CryoSPARC^39^, which reconstructed a trajectory for PRIM2_CTD_ moving from the initiation position away from PRIM1(Figure 4b, Movie S3). This evidence supports the notion that the PRIM2_CTD_ remains engaged with the 5’-end of the growing RNA chain during primer synthesis.

**Figure 4.**
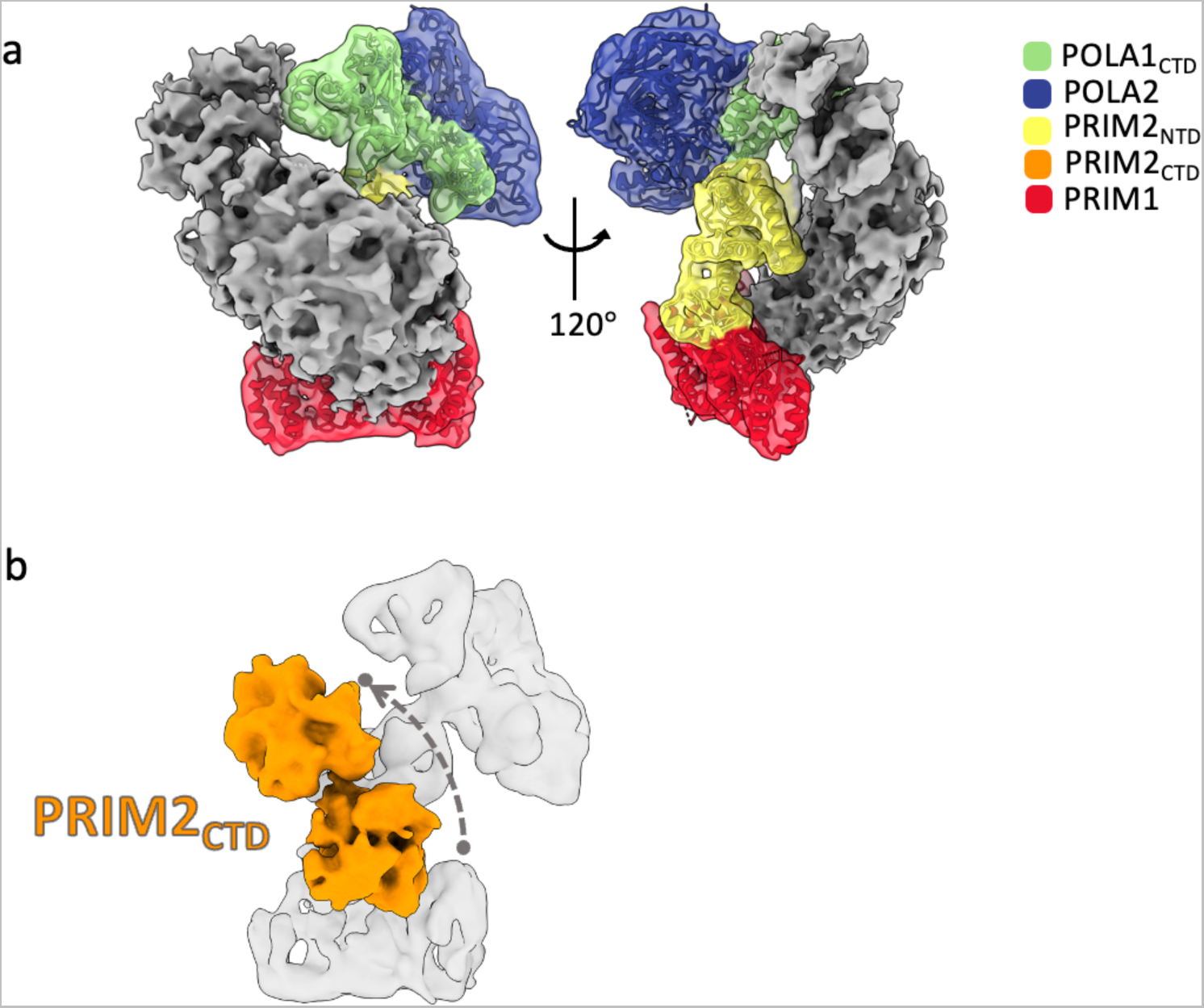
PRIM2_CTD_ tracks the 5’-end of the growing RNA primer. (**a**) CryoEM map of primosome from priming reactions with ATP only. Contouring at a lower threshold (σ=0.064) reveals the dynamic position of PRIM2_CTD_, compatible with its trajectory during primer synthesis. PRIM2_CTD_ density is shown as solid grey surface. The platform components are fitted in the transparent density and coloured as in Figure 1. (**b**) Volume series obtained from the 3D variability analysis, showing the start and end positions of PRIM2_CTD_ in the series.

### RNA primer termination requires POLA1 polymerase domain and deoxynucleotide

Our cryoEM analysis so far had provided insights into the conformational changes that the primosome undergoes to initiate and extend the RNA primer. We sought to analyse the subsequent step of RNA primer termination by increasing the incubation time of the priming reaction, in the presence of both ATP and dATP, before sample freezing and cryoEM data collection (Figure S1b, Table S1, Figure S8). We identified a population of primosome particles where density attributable to the POLA1 polymerase domain – that had become mobile at initiation following its release from the primosome platform– was visible in the map (Figure 5a). Although the resolution of the map is insufficient for their accurate relative positioning, POLA1 polymerase domain is in contact with the PRIM2_CTD_ bound to the 5’-end of the primer, in agreement with a role for POLA1 in RNA primer termination. Thus, blocking the path of the PRIM2_CTD_ tracking the growing primer would be part of the mechanism that POLA1 adopts to trigger primer release from PRIM1 and take over primer elongation with deoxynucleotides.

**Figure 5.**
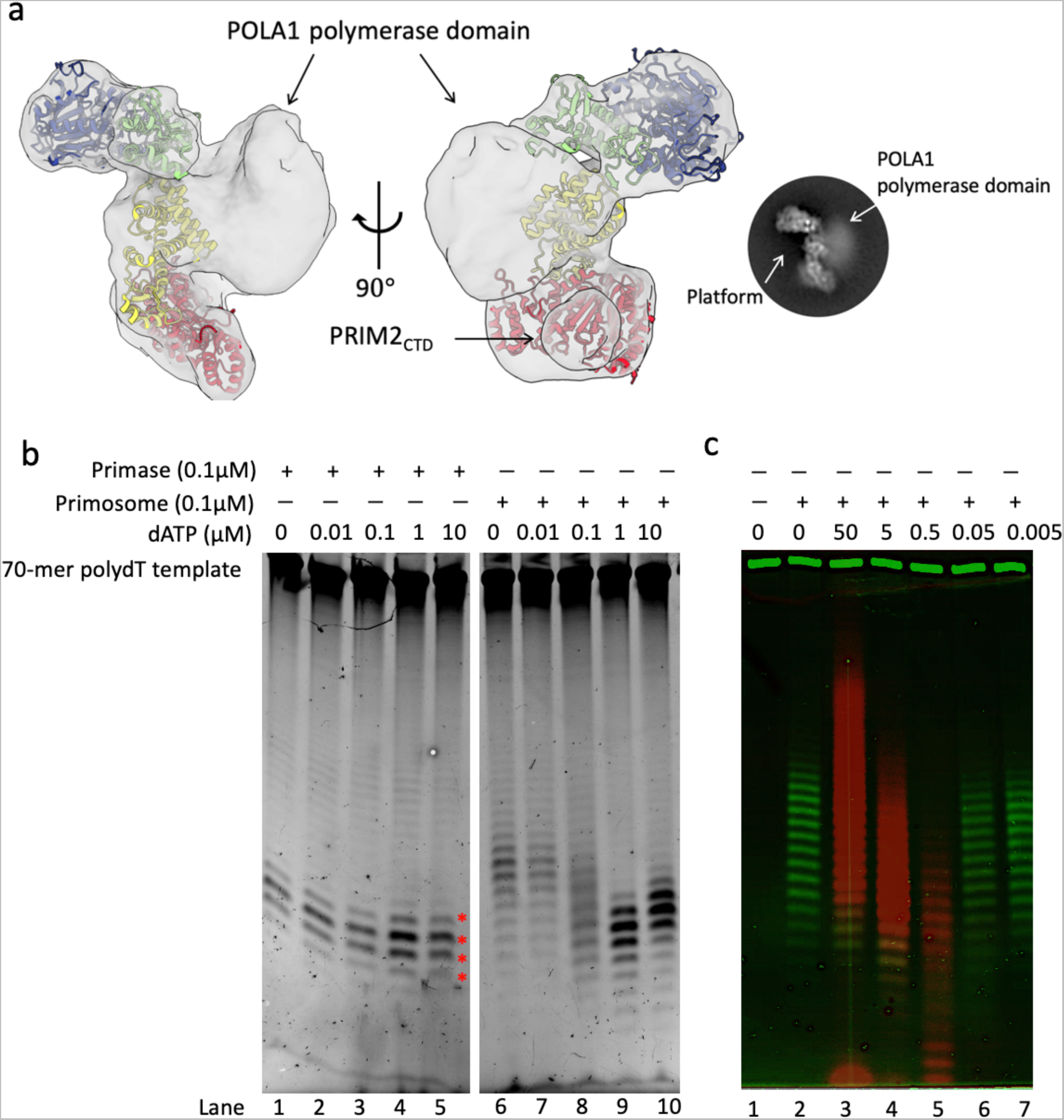
Role of POLA1 in specification of RNA primer size. (**a**) Two views of the cryoEM map of the primosome showing the POLA1 polymerase domain and PRIM2_CTD_ in a reciprocal position compatible with RNA synthesis termination. A model of the primosome platform from initiation state has been rigid-body fitted into the map. The oval shows a representative 2D class highlighting the density of POLA1 polymerase domain. (**b**) Urea-PAGE analysis of dATP titration in priming reactions with primase or primosome. Lanes 1-5, primase activity; lanes 6-10, primosome activity. The red stars mark the RNA products in size range from 9 to 12 nucleotides. (**c**) Urea-PAGE analysis of Cy5-dATP (red) titration in priming reactions of the primosome. Nucleic acid products were stained with Sybr gold (green).

To further investigate the role of POLA1 in regulating RNA primer synthesis, we titrated increasing concentration of dATP in the primer synthesis reaction by the primosome. While RNA primer synthesis is not affected by dATP in a control reaction using primase, it is clearly modulated by dATP in the context of the primosome: primer size distribution becomes sharper and centred around a shorter mean size of 9 nucleotides (Figure 5b). To be able to track DNA incorporation into the primer, we used Cy5-labelled dATP (Figure 5c). The experiment makes it evident that POLA1 has a direct dNTP-dependent role in determining primer size: while the primase activity of the primosome produces longer RNA primers with a wide range of sizes, POLA1 and dATP combine to capping RNA size to 9 - 12 nucleotides.

Taken together, the evidence from our cryoEM and biochemical experiments indicates that POLA1 contributes to determine the size of the RNA primer and that this effect is dependent on dATP. Thus – in the absence of or at insufficient concentrations of dATP – the polymerase domain of POLA1 is unable to position itself to intercept the moving PRIM2_CTD_, leaving PRIM1 free to continue RNA polymerisation.

## DISCUSSION

In this study, we have visualised conformational states of the human primosome during RNA primer synthesis using cryoEM snapshots of the priming reaction. Our approach is complementary to previous studies that have focused on the mechanism of primer elongation by providing the primosome with an RNA-primed DNA template substrate^29, 40^. Taken together, our structural evidence can be combined in a mechanism of RNA primer synthesis by the human primosome (Figure 6).

**Figure 6.**
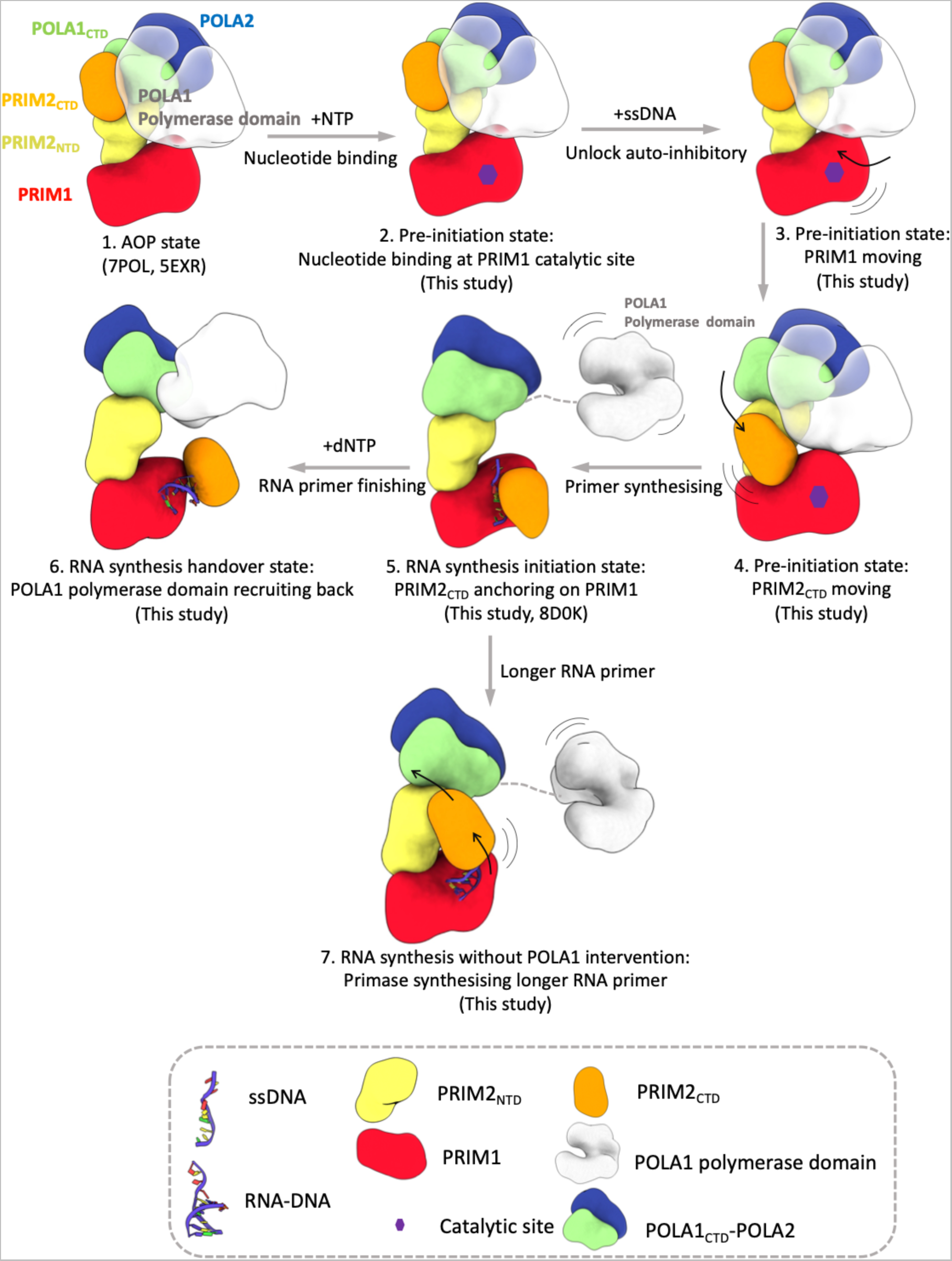
Model of RNA primer synthesis by the human primosome. 1. Apo state of the human primosome complex. 2. Nucleotide loading of PRIM1. 3. Unlocking of PRIM1 from its anchoring interaction with POLA1 polymerase domain and movement towards PRIM2_CTD_. 4. PRIM2_CTD_ moves towards PRIM1 and docks on its active site. 5. Release of POLA1 polymerase domain from the platform licences PRIM1 and PRIM2_CTD_ to start RNA primer synthesis. 6. POLA1 polymerase domain is recruited back to the platform and engages PRIM2_CTD_ to promote RNA primer handover. 7. In the absence of dATP, POLA1 polymerase domain is unable to effect primer handover, resulting in continued primer synthesis by primase.

### Species-dependent regulation of the apo state

The crystal structure of the human primosome in the apo state revealed a compact arrangement of subunits^25^, where PRIM1, POLA1 polymerase domain and PRIM2_CTD_ are sequestered in unproductive interactions with a structural platform formed by PRIM1-PRIM2_NTD_ - POLA1_CTD_ - POLA2. Later cryoEM studies of the human primosome bound to the SARS-CoV-2 nsp1 protein^26^ and to the telomeric CST complex confirmed the existence of an apo state^30^. Interestingly, structural studies of primosome from other eukaryotic species have revealed that the apo state is not as strictly maintained as in the human complex. Thus, the primosome from *Xenopus laevis* shows a mixture of apo and initiation states^29^, while in the apo primosome of *S. cerevisiae* PRIM1 is disengaged from POLA1 and available for interactions with the DNA template^28^. In our experiments with the human primosome we have shown that the provision of ssDNA alone is insufficient to induce significant conformational changes in the primosome. Furthermore, we found that the human primosome in pre-initiation stages shares similar feature of a flexible PRIM1 as the apo state of primosome in *S. cerevisiae*^28^. Amino acid differences in the sequence of PRIM1 anchor loop (Figure S9) might explain the looser anchoring of yeast PRIM1. Moreover, we have identified a distinct conformation of the apo state with nucleotide bound in PRIM1’s active site, under reaction conditions that include ssDNA, ATP and dATP, suggesting that the first stage of initiation involves nucleotide binding to PRIM1 without concurrent conformational alterations.

Collectively, these findings indicate that ‘apo’ primosomes from different species maintain an auto-inhibitory conformation with varying degree of stringency. A tighter regulation of the human primosome might be required in DNA replication. Alternatively, it might be related to the proposed non-replicative role of Pol α - primase in the synthesis of cytosolic RNA: DNA primers, to modulate the innate immune system by dampening the interferon I response^41^.

### From the apo state to the initiation state

Initiation of RNA primer synthesis requires transition of the primosome from the apo state to pre-initiation state where the primase subunits adopt the correct reciprocal arrangement for catalysis. Our study identifies four intermediate states which shed light on the primosome trajectory to the initiation state. The states show concerted rigid-body movements of PRIM1 and the PRIM2_CTD_ towards each other, that detach PRIM2_CTD_ from POLA1’s polymerase domain and brings it on top of PRIM1’s active site, while the rest of the primosome maintains its overall apo conformation. This reciprocal approach is followed by an in-place rotation of PRIM2_CTD_ that brings its NTP- and DNA-binding residues in line with PRIM1’s catalytic residues for the initiation reaction. The flexible linker that connects PRIM2_CTD_ to PRIM2_NTD_ must be essential for this rearrangement, as well as for PRIM2_CTD_’s ability to remain bound to the 5’-end of the primer during primer extension^12^.

The primosome conformations at the pre-initiation stage 4 and initiation state share a common arrangement of the platform and PRIM2_CTD_. However, they differ in that the polymerase domain of POLA1 remains attached to the platform in pre-initiation stage, whereas it has become mobile in the initiation state. This difference suggests that the release of POLA1 polymerase domain is required for primer synthesis to begin, possibly by allowing PRIM1 and PRIM2_CTD_ to achieve the optimal reciprocal positioning for initiation of primer synthesis.

In the study of human primosome - CST complex bound to telomeric DNA^32^, the PRIM2_CTD_ was positioned in front of PRIM1, while the DNA template was threaded between the primase subunits. The initiation stage of the primosome in our study shows a similar orientation of PRIM1 and PRIM2_CTD_ (Figure S6). Without the conformational restraints posed by the CST on primase and the DNA template, PRIM2_CTD_ remains highly mobile, and we cannot resolve the template DNA in our map. The single-stranded DNA binding protein RPA is known to interact with Pol α - primase and it is possible that it assists priming in a similar fashion to CST in DNA replication^42^.

### Primer elongation and hand-off to POLA1

According to current models of primer synthesis^28, 29, 43^, the PRIM2_CTD_ remains bound to the 5’- end of the primer during primer elongation, thanks to its flexible tethering to the PRIM2_NTD_. By applying 3D variability analysis to our cryoEM study of the priming reaction, we have been able to reconstruct a trajectory for PRIM2_CTD_ that shows its movement away from PRIM1 as it tracks the 5’-end of the growing primer (Movie S3). Thus, our structural analysis of the active human primosome provides experimental evidence that supports the continuous engagement of PRIM2_CTD_ with the RNA primer during synthesis.

Our biochemical data show that POLA1 has a role in shaping primer synthesis, as primer size and length distribution are both clearly modulated by it. This observation implies an active mechanism of primer hand-over whereby POLA1 takes over primer synthesis. Our data further show that primer takeover by POLA1 requires dNTP, indicating that the deoxynucleotide is important for the polymerase to take up the correct conformation for primer extension. Consistent with our biochemical data, the recruitment of POLA1 was observed at the elongation stage of RNA primer synthesis (Figure 5a). The recent cryoEM study of primosome in. *S. cerevisiae* has shown a similar intermediate configuration where the polymerase domain of POLA1 is hanging above PRIM2 in preparation for handover of the mature RNA primer^28^. Taken together, this evidence indicates that the POLA1 polymerase domain controls the size of RNA primer.

Thus, our findings have revealed an intimate involvement of the polymerase domain of POLA1 at all stages of primer synthesis: as the primosome transitions towards the initiation state, the polymerase domain remains in contact with the primase subunits until a late pre-initiation stage, when it must be released from the platform for initiation to begin; it is then recruited back to the platform to terminate RNA synthesis, thus contributing to specify primer size.

In summary, we have described experimental evidence that sheds light on the sequential transformations of the human primosome from its initial inactive state to the initiation of RNA primer synthesis, primer elongation and hand over to POLA1. This information, together with existing complementary evidence on primer elongation by the primosome, provides a structural basis for a complete description of the mechanism of RNA - DNA primer synthesis by the human primosome.

### Data Availability

The cryo-EM density maps have been deposited in the Electron Microscopy Data Bank (https://www.ebi.ac.uk/pdbe/emdb) with accession numbers EMD-17795 (pre-initiation stage 1); EMD-17807 (pre-initiation stage 2); EMD-17810 (pre-initiation stage 3); EMD-17811 (pre-initiation stage4); EMD-17812 (initiation I); EMD-17813 (initiation II); EMD-17824 (handover).

## Acknowledgements

We would like to thank Dimitri Chirgadze and staff at the cryoEM Facility of the Department of Biochemistry for help with data collection. This work was funded by Wellcome Trust award 221892/Z/20/Z to L.P..

