## Supplemental Figures for "CryoEM insights into RNA primer synthesis by the human primosome"

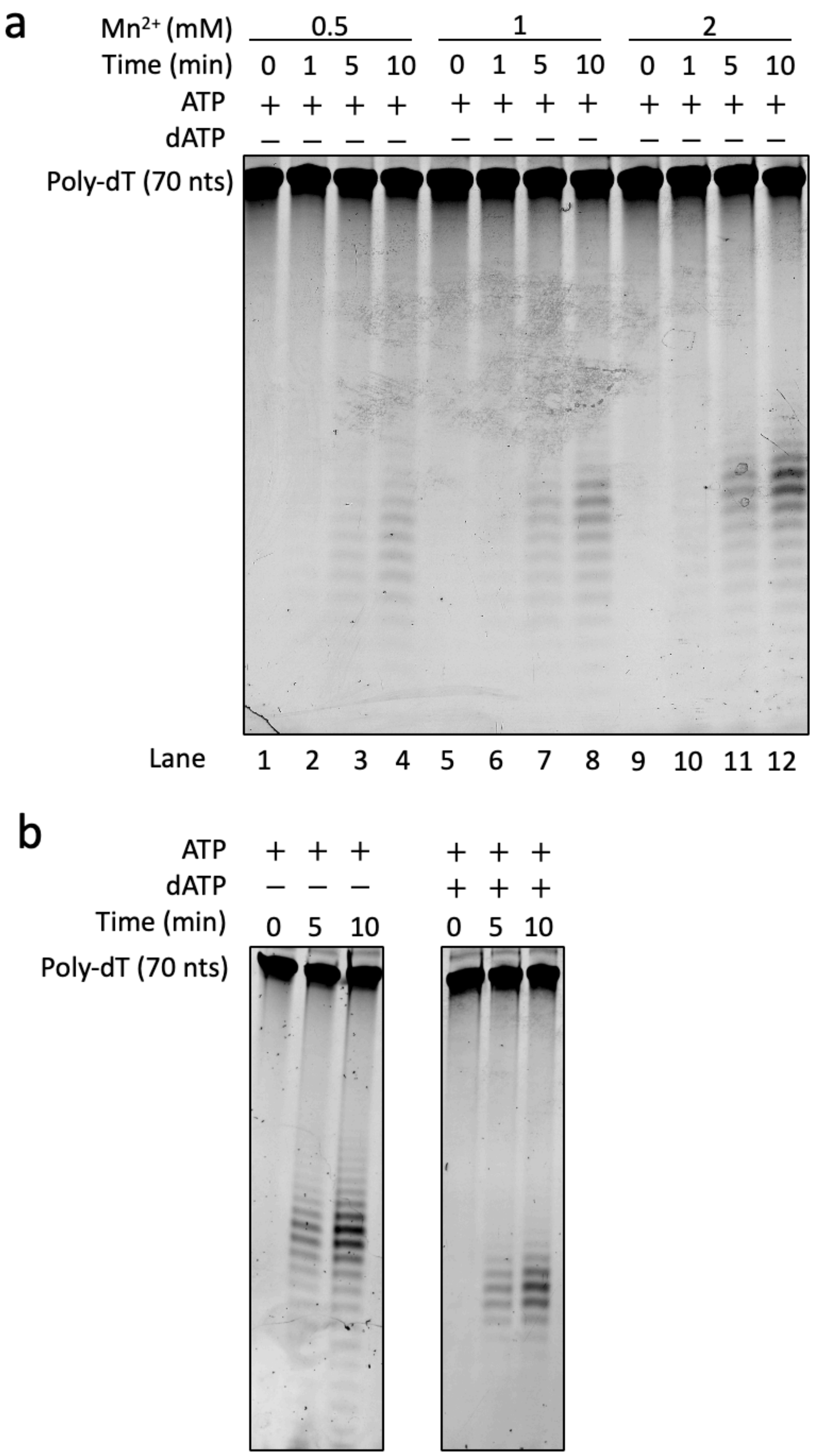
a
Mn2+ (mM)
0.5
1
2
Time (min)
0 1 5 10 0 1 5 10 0 1 5 10
ATP
+ + + + + + + + + + + +
dATP
– – – – – – – – – – – –
Poly-dT (70 nts)
Lane
1 2 3 4 5 6 7 8 9 10 11 12
b
ATP
+ + + + + +
dATP
– – – + + +
Time (min)
0 5 10 0 5 10
Poly-dT (70 nts)

**Figure S1.** Optimisation of primosome activity for cryoEM analysis using denaturing urea-PAGE. Primosome at 0.1 μM and poly-dT 70mer at 5 μM were used together with the indicated amounts of $Mn^{2+}$, ATP and dATP. (**a**) Time course of priming reaction in the presence of 0.5 mM ATP and increasing concentration of manganese. (**b**) Optimisation of reaction time for RNA primer initiation (0.5 mM ATP only, left panel) and handover (0.5 mM ATP and 0.05 mM dATP, right panel). The reaction buffer contains 2 mM manganese as the optimised concentration from (a).

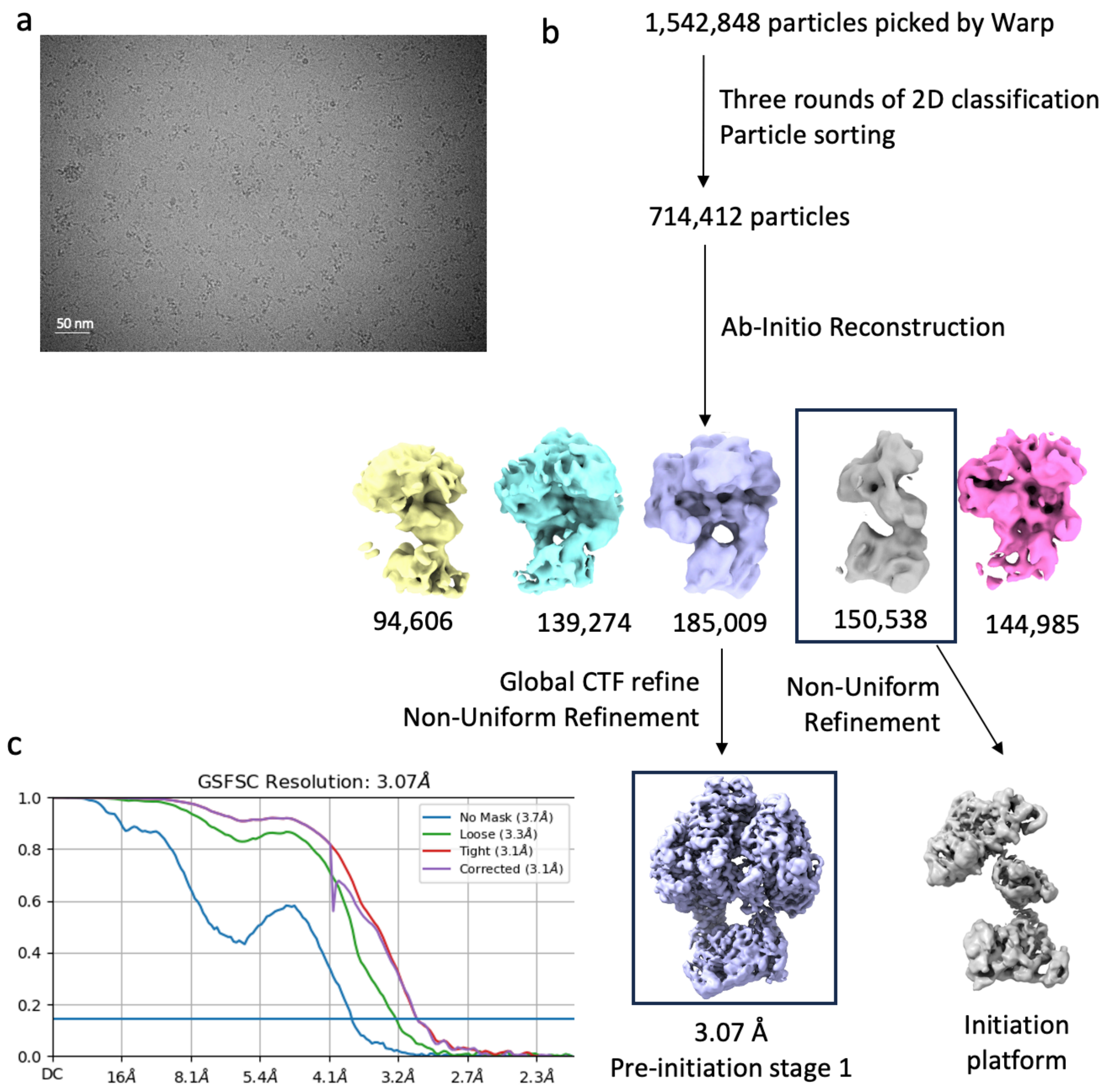


**Figure S2.** Cryo-EM data processing of the human primosome from a priming reaction with polydT-ssDNA 70mer, ATP and dATP (Table S1 – pre-initiation stage 1). (**a**) Representative motion-corrected micrograph. (**b**) Cryo-EM image-processing pipeline used for primosome initiation, including particle classification, selection, and 3D refinement on the selected 3D class. (**c**) The global resolution estimate from the masked Fourier Shell Correlation curve is 3.07 Å at FSC of 0.143 for the map of pre-initiation stage 1.

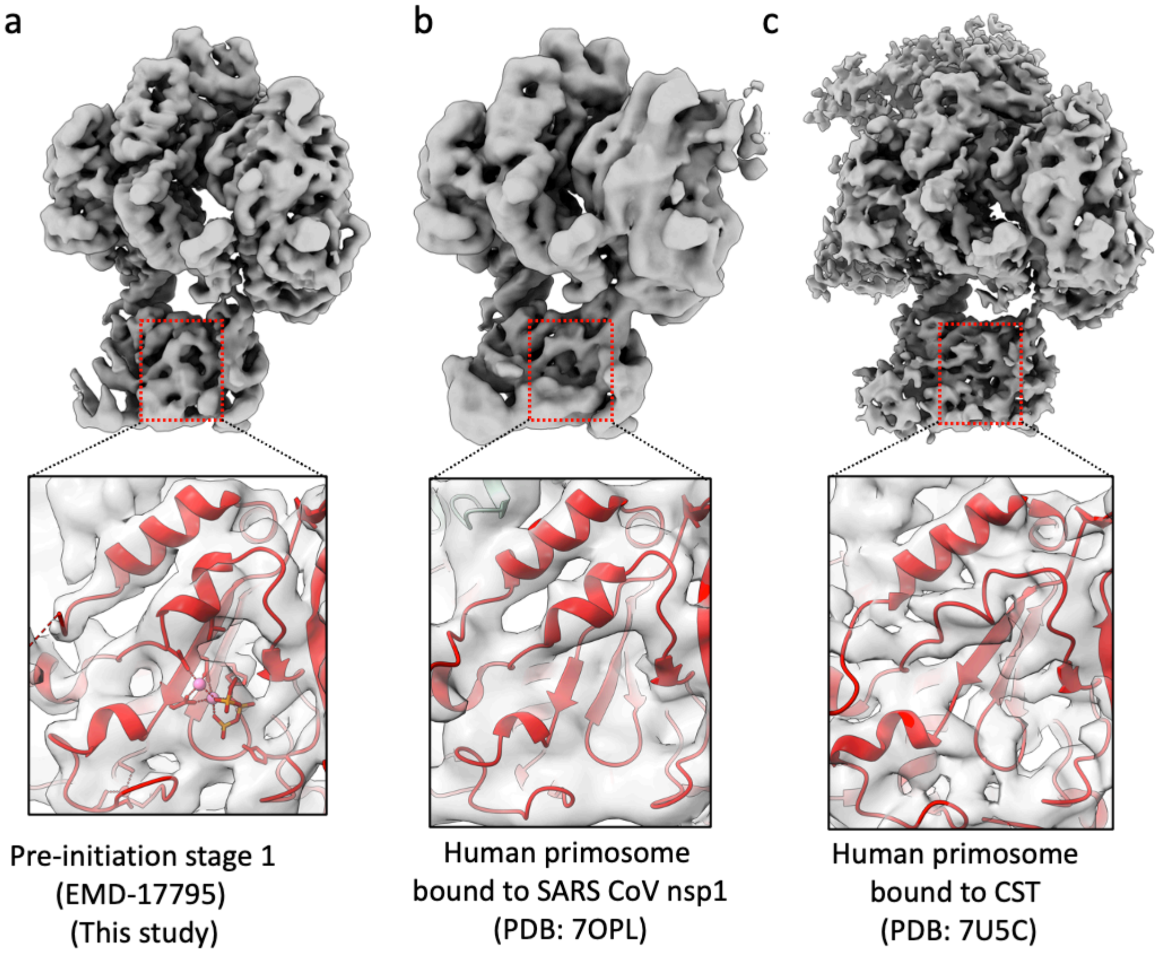


**Figure S3.** Comparison of PRIM1 active site in cryoEM maps of apo-state of the human primosome. Details of the PRIM1 density are shown in the box below each map. In (a) the PRIM1 structure (PDB:6R4S) is rigid-body fitted into the map. (**b**) The model of human primosome in apo state bound to nsp1 (PDB: 7OPL) in cryoEM density map (EMD-13020). (**c**) The model of human primosome in recruitment/apo state bound to CST (PDB:7U5C) in the cryoEM density map (EMD-26346).

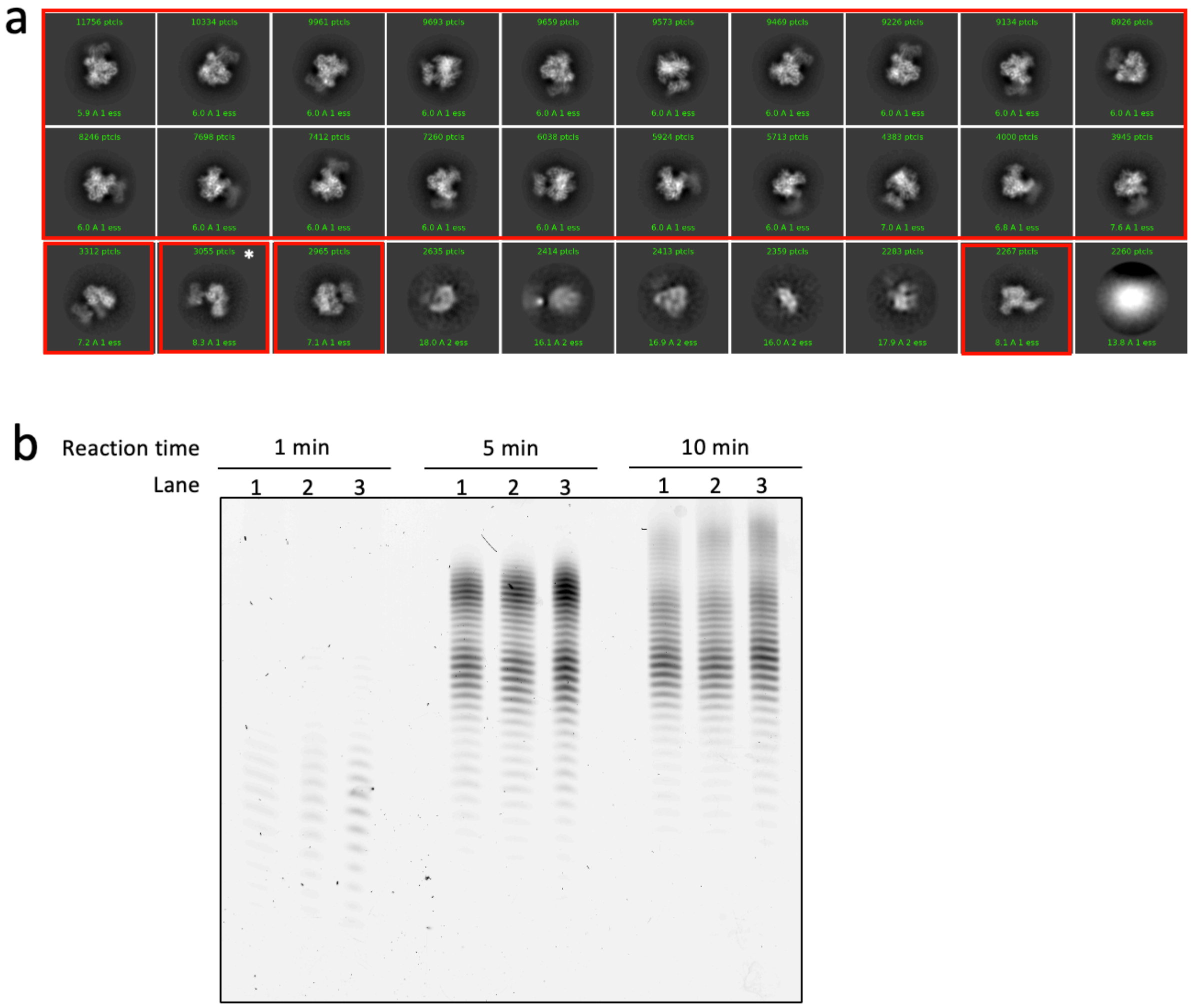


| Lane | Pre-incubation conditions |
|---|---|
| 1 | No pre-incubation |
| 2 | Primosome pre-incubates with ssDNA template Poly-dT (70 nts) |
| 3 | Primosome pre-incubates with nucleotide (ATP) |

**Figure S4.** CryoEM and urea-PAGE assay of priming by the primosome with either ssDNA or ATP pre-incubation. (**a**) CryoEM 2D classification of primosome particles from a priming reaction that had been pre-incubated with ssDNA overnight. Top 25 classes are shown ranked by number of particles. Selected classes are shown in red boxes. Asterisks (*) marks the only class with unassigned conformation. 98.2% of the 168,946 selected particles are in the apo conformation. (**b**) Denaturing urea PAGE assay of time-course priming reaction at 20 °C with 0.1 µM primosome pre-incubation at 20 °C for 15 mins with either 5 µM ssDNA or 0.5 mM ATP.

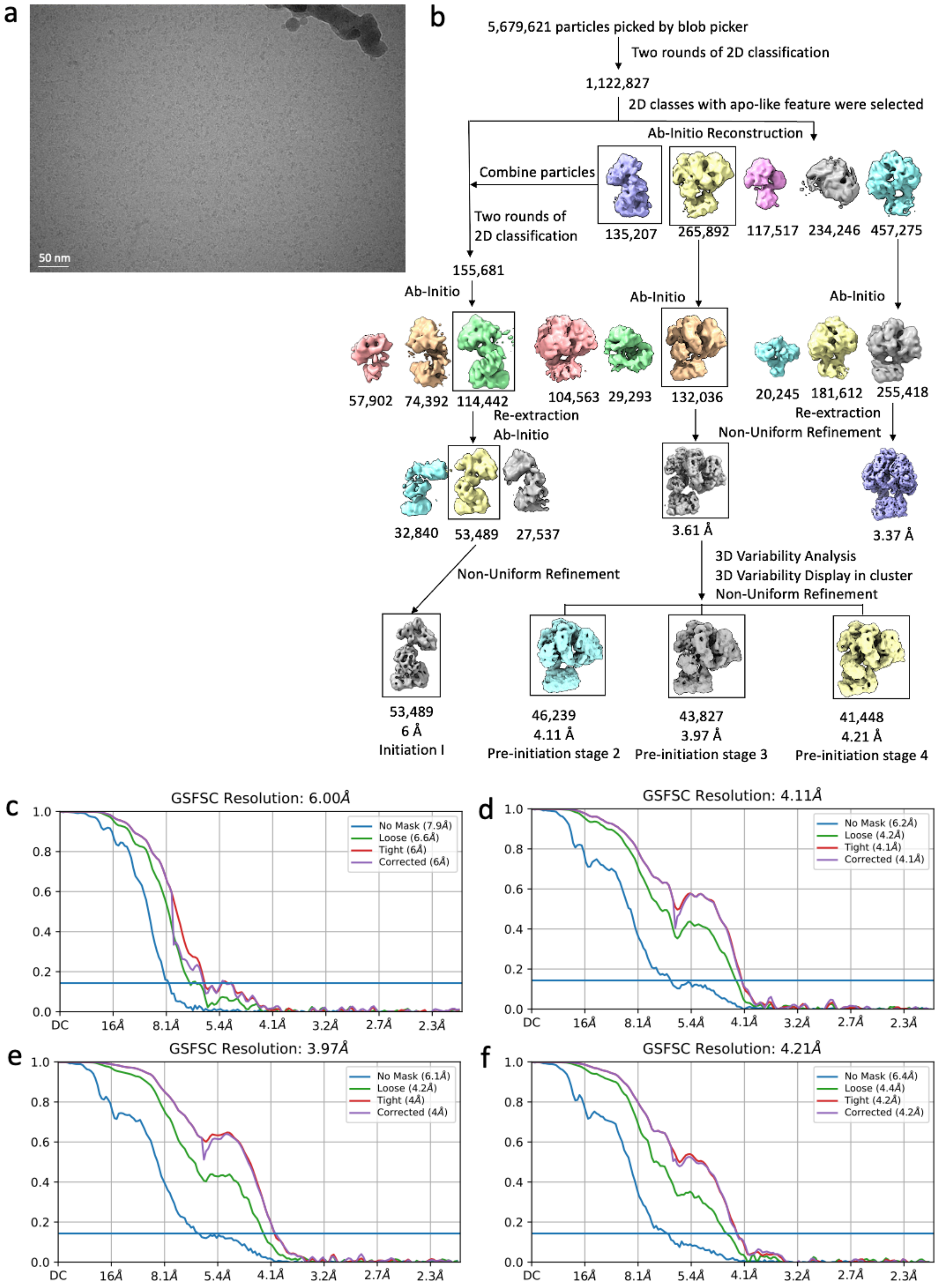
a
50 nm
b
5,679,621 particles picked by blob picker
Two rounds of 2D classification
1,122,827
2D classes with apo-like feature were selected
Ab-Initio Reconstruction
Combine particles
Two rounds of 2D classification
135,207
265,892
117,517
234,246
457,275
155,681
Ab-Initio
Ab-Initio
Ab-Initio
57,902
74,392
114,442
104,563
29,293
132,036
20,245
181,612
255,418
Re-extraction
Ab-Initio
Re-extraction
Non-Uniform Refinement
32,840
53,489
27,537
3.61 Å
3.37 Å
Non-Uniform Refinement
3D Variability Analysis
3D Variability Display in cluster
Non-Uniform Refinement
53,489
6 Å
Initiation I
46,239
4.11 Å
Pre-initiation stage 2
43,827
3.97 Å
Pre-initiation stage 3
41,448
4.21 Å
Pre-initiation stage 4
c
GSFSC Resolution: 6.00Å
No Mask (7.9Å)
Loose (6.6Å)
Tight (6Å)
Corrected (6Å)
d
GSFSC Resolution: 4.11Å
No Mask (6.2Å)
Loose (4.2Å)
Tight (4.1Å)
Corrected (4.1Å)
e
GSFSC Resolution: 3.97Å
No Mask (6.1Å)
Loose (4.2Å)
Tight (4Å)
Corrected (4Å)
f
GSFSC Resolution: 4.21Å
No Mask (6.4Å)
Loose (4.4Å)
Tight (4.2Å)
Corrected (4.2Å)
DC
16Å
8.1Å
5.4Å
4.1Å
3.2Å
2.7Å
2.3Å

**Figure S5.** CryoEM data processing scheme of the human primosome from a priming reaction with polydT-ssDNA 70mer, ATP and dATP with $BS^3$ (Table S1 – pre-initiation stage 2,3,4, and Initiation state I) (**a**) Representative motion-corrected micrograph. (**b**) Cryo-EM image-processing pipeline showing particle classification, selection, and 3D refinement on selected 3D classes. In this pipeline, 3 ab-initio models with distinct features were selected and went through another ab-initio to reduce heterogeneity of the classes before 3D refinement was performed. (**c**), (**d**), (**e**) and (**f**) shows the global resolution estimation of the pre-initiation stage 2,3,4, and Initiation state I respectively, using the gold standard FSC and a cut-off criterion of 0.143.

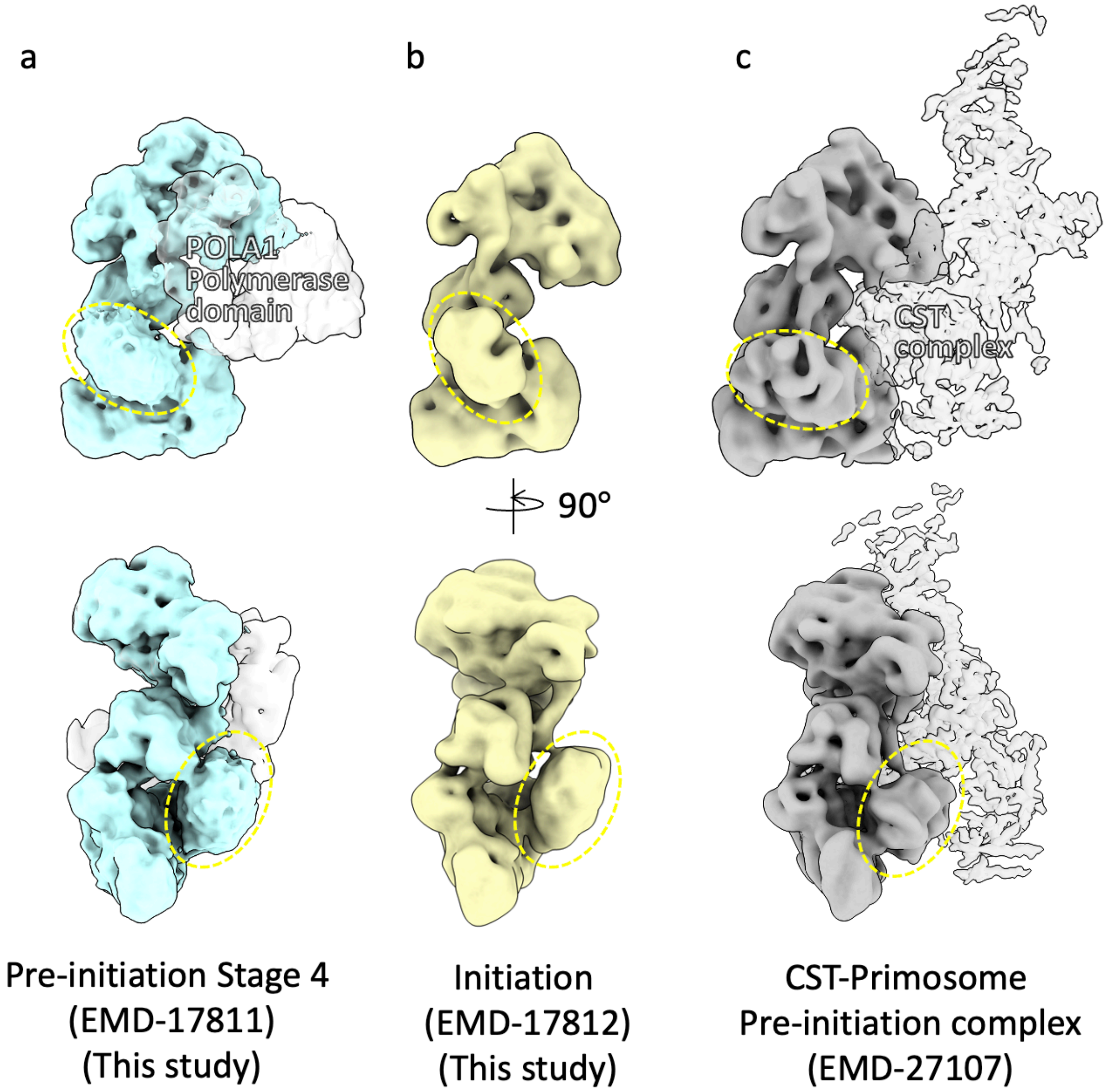


**Figure S6**. PRIM2$_{CTD}$ position in the cryoEM maps of the primosome at the pre-initiation (**a**, **c**) and initiation (**b**) states of primer synthesis. Two rotated views for each map are shown. The density of the primosome platform (PRIM1-PRIM2$_{NTD}$-POLA1$_{CTD}$-POLA2) is shown as a solid surface and the PRIM2$_{CTD}$ density is highlighted by a yellow dashed oval. The densities for POLA1 polymerase domain (a) and the CST, DNA and POLA1 (c) are shown as transparent surfaces. In (c), the map of the human CST - Pol $\alpha$ - primase preinitiation complex was low-pass filtered to 6 Å.

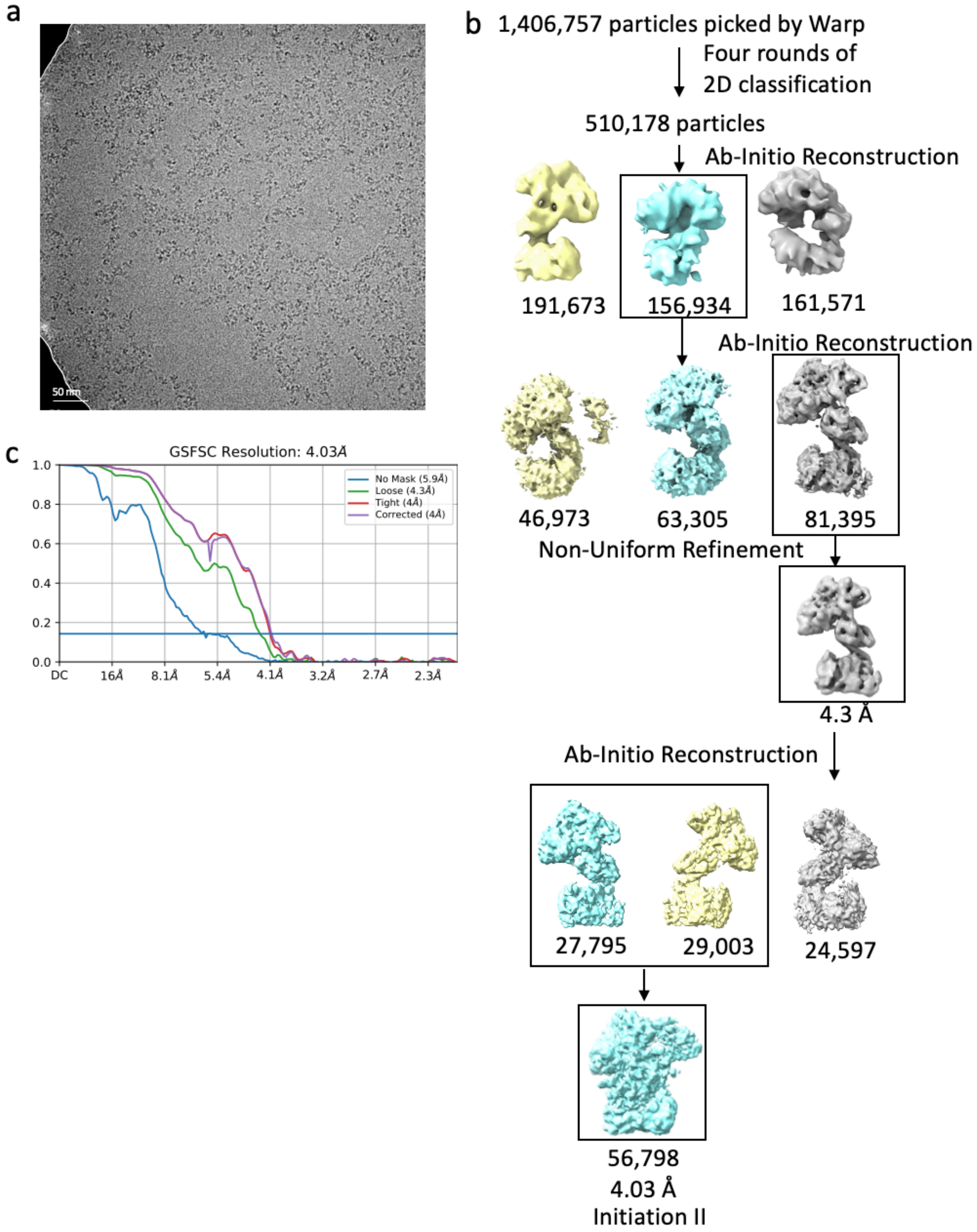


**Figure S7.** CryoEM data processing scheme of human primosome from a priming reaction with polydT-ssDNA 70mer and ATP (Table S1 –Initiation state II). (**a**) An example of the screening micrograph. (**b**) Cryo-EM image-processing pipeline showing particle classification, selection, and 3D refinement on selected 3D classes. The map of initiation state was then gone through 3D variability analysis. (**c**) The global resolution estimate from the masked Fourier Shell Correlation curve is 4.03 Å at FSC = 0.143 for initiation state.

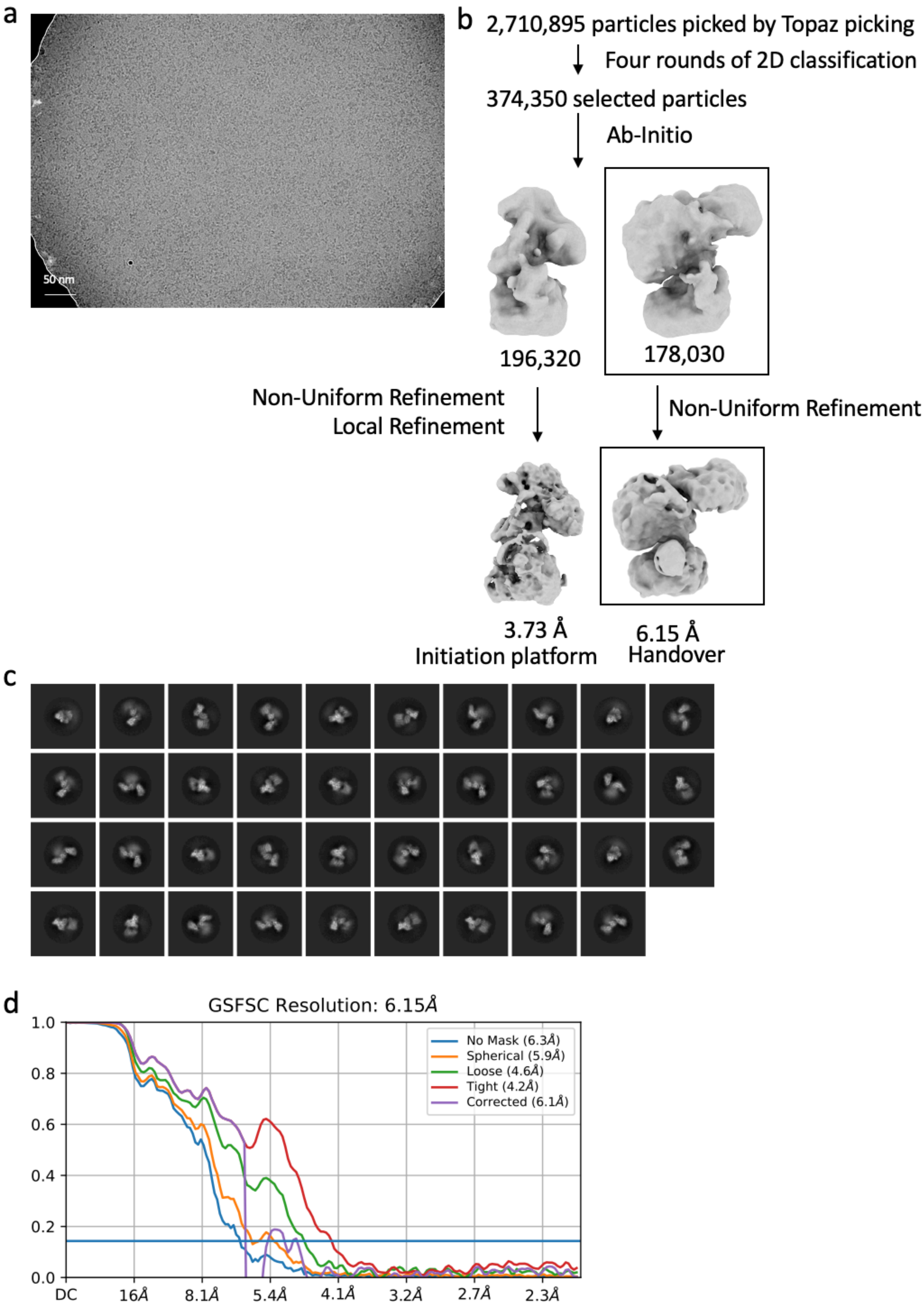
a
50 nm
b
2,710,895 particles picked by Topaz picking
Four rounds of 2D classification
374,350 selected particles
Ab-Initio
196,320
178,030
Non-Uniform Refinement
Local Refinement
Non-Uniform Refinement
3.73 Å
Initiation platform
6.15 Å
Handover
c
d
GSFSC Resolution: 6.15Å
No Mask (6.3Å)
Spherical (5.9Å)
Loose (4.6Å)
Tight (4.2Å)
Corrected (6.1Å)
1.0
0.8
0.6
0.4
0.2
0.0
DC
16Å
8.1Å
5.4Å
4.1Å
3.2Å
2.7Å
2.3Å

**Figure S8.** CryoEM data processing scheme of human primosome from a longer priming reaction with polydT-ssDNA 70mer, ATP and dATP with $BS^3$. (**a**) Representative motion-corrected micrograph. (**b**) Cryo-EM image-processing pipeline showing particle classification, selection, and 3D refinement on selected 3D classes. At the beginning of the pipeline, Topaz trained with reference particles showing initiation state in 2D class was used to select desired particles. (**c**) Representative 2D-class averages show high-resolution features and different orientations. (**d)** The global resolution estimate from the masked Fourier Shell Correlation curve is 6.15 Å at FSC of 0.143 for the handover state.

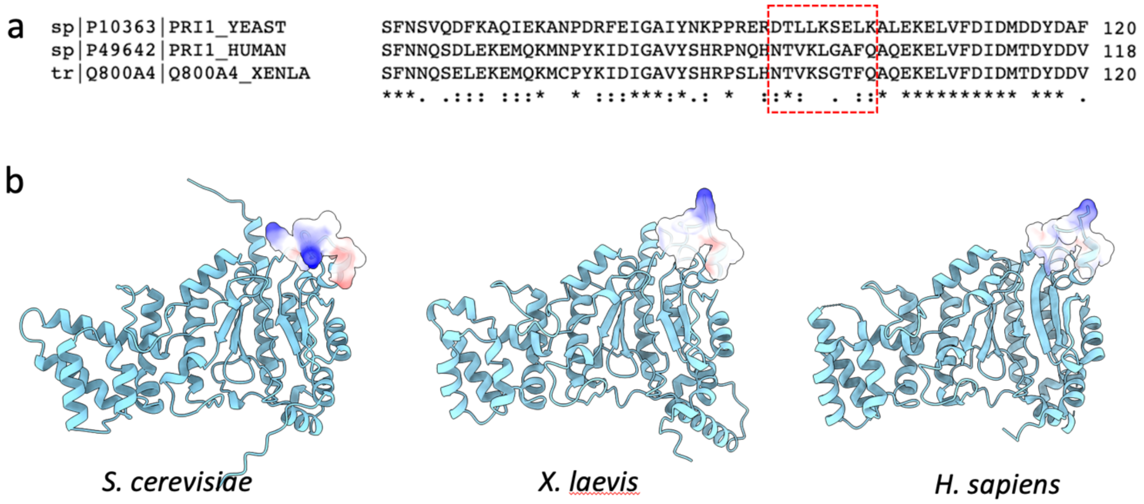


**Figure S9.** Comparison of primase small subunit between species. (**a**) Multiple sequence alignment of PRIM1 performed using Clustal-O (1.2.4). The dashed red square shows the sequence of the anchor loop N92-Q100. Asterisks (*) indicate positions with identical residues, colons (:) and periods (.) indicate strongly and weakly conserved residues. (**b**) The electrostatic charge of amino acids in loop N92-Q100 is shown for yeast, frog, and human PRIM1, as molecular surface coloured red (negative) to blue (positive). The models of *S. cerevisiae* and *X. laevis* PRIM1 were generated by AlphaFold structure prediction; the model of *H. sapiens* PRIM1 was obtained from PDB:7OPL.

### Table S1. Cryo-EM data collection and processing

| **Data set** | Pre-initiation | | | | Initiation | | Handover |
|---|---|---|---|---|---|---|---|
| | stage 1 | stage 2 | stage 3 | stage 4 | I | II | |
| **Reaction condition** | Primosome<br>+ ssDNA<br>+ ATP+ dATP<br>5 mins at 20°C | Primosome<br>+ ssDNA<br>+ ATP + dATP<br>5 mins at 20°C<br>($BS^3$ cross-linked) | | | | Primosome<br>+ ssDNA<br>+ ATP<br>5 mins at 20°C | Primosome<br>+ ssDNA<br>+ ATP + dATP<br>10 mins at 20°C<br>($BS^3$ cross-linked) |
| EMDB code | EMD-17795 | EMD-17807 | EMD-17810 | EMD-17811 | EMD-17812 | EMD-17813 | EMD-17824 |
| **Data collection** | | | | | | | |
| Voltage (kV) | 300 | 300 | | | | 300 | 300 |
| Magnification (nominal) | 81,000 | 81,000 | | | | 81,000 | 81,000 |
| Calibrated Pixel size (Å) | 1.066 | 1.066 | | | | 1.066 | 1.066 |
| Symmetry imposed | C1 | C1 | | | | C1 | C1 |
| Defocus range (μm) | -1.0 to -2.6 | -1.0 to -2.6 | | | | -1.0 to -2.6 | -1.0 to -2.6 |
| Electron exposure ($e^-/Å^2$) | 47.78 | 50.50 | | | | 46.05 | 53.67 |
| Number of Fractions | 45 | 45 | | | | 40 | 45 |
| Final images (no.) | 15,430 | 7,014 | | | | 10,677 | 14,621 |
| **Data processing** | | | | | | | |
| Software | Warp,<br>cryoSPARC<br>(4.0.0) | cryoSPARC (4.2.1) | | | | Warp,<br>cryoSPARC<br>(v3.3.2) | cryoSPARC<br>(4.2.1) |
| Particle number | 185,009 | 46,239 | 43,827 | 41,448 | 53,489 | 56,798 | 178,030 |
| Map sharpening B factor ($Å^2$) | -85.3 | -130.4 | -126.5 | -139.6 | -448.6 | -126.4 | -252.6 |
| Map resolution (Å)<br>(FSC=0.143) | 3.07 | 4.11 | 3.97 | 4.21 | 6.0 | 4.03 | 6.15 |

**Movie S1.** Volume series movie generated from 3D variability analysis of the primosome map showing the transition from preinitiation stage 2 to 4.

**Movie S2.** Model of the primosome platform morphing between apo and initiation states. Colours of the subunits are the same as in Figure 1.

**Movie S3.** Volume series movie generated from 3D variability analysis of the primosome map showing the movement of $PRIM2_{CTD}$ during RNA primer synthesis.
